# Freshly Thawed Cryobanked Human Neural Stem Cells Engraft within Endogenous Neurogenic Niches and Restore Cognitive Function Following Chronic Traumatic Brain Injury

**DOI:** 10.1101/2020.07.23.212423

**Authors:** Anna Badner, Emily K. Reinhardt, Theodore V. Nguyen, Nicole Midani, Andrew T. Marshall, Cherie Lepe, Karla Echeverria, Javier Lepe, Vincent Torrecampo, Sara H. Bertan, Serinee H. Tran, Aileen J. Anderson, Brian J. Cummings

## Abstract

Human neural stem cells (hNSCs) have potential as a cell therapy following traumatic brain injury (TBI). While various studies have demonstrated the efficacy of NSCs from on-going culture, there is a significant gap in our understanding of freshly thawed cells from cryobanked stocks – a more clinically-relevant source. To address these shortfalls, the therapeutic potential of our previously validated Shef-6.0 human embryonic stem cell (hESC)-derived hNSC line was tested following long-term cryostorage and thawing prior to transplant. Immunodeficient athymic nude rats received a moderate unilateral controlled cortical impact (CCI) injury. At 4-weeks post-injury, 6×10^5^ freshly thawed hNSCs were transplanted into six injection sites (2 ipsi- and 4 contra-lateral) with 53.4% of cells surviving three months post-transplant. Interestingly, most hNSCs were engrafted in the meninges and the lining of lateral ventricles, associated with high CXCR4 expression and a chemotactic response to SDF1alpha (CXCL12). While some expressed markers of neuron, astrocyte, and oligodendrocyte lineages, the majority remained progenitors, identified through doublecortin expression (78.1%). Importantly, transplantation resulted in improved spatial learning and memory in Morris water maze navigation and reduced risk-taking behavior in an elevated plus maze. Investigating potential mechanisms of action, we identified an increase in ipsilateral host hippocampus cornu ammonis (CA) neuron survival, contralateral dentate gyrus (DG) volume and DG neural progenitor morphology as well as a reduction in neuroinflammation. Together, these findings validate the potential of hNSCs to restore function after TBI and demonstrate that long-term bio-banking of cells and thawing aliquots prior to use may be suitable for clinical deployment.

**Significance Statement:** There is no cure for chronic traumatic brain injury (TBI). While human neural stem cells (hNSCs) offer a potential treatment, no one has demonstrated efficacy of thawed hNSCs from long-term cryobanked stocks. Frozen aliquots are critical for multisite clinical trials, as this omission impacted the use of MSCs for graft versus host disease. This is the first study to demonstrate the efficacy of thawed hNSCs, while also providing support for novel mechanisms of action – linking meningeal and ventricular engraftment to reduced neuroinflammation and improved hippocampal neurogenesis. Importantly, these changes also led to clinically relevant effects on spatial learning/memory and risk-taking behavior. Together, this new understanding of hNSCs lays a foundation for future work and improved opportunities for patient care.

## Introduction

Traumatic brain injury (TBI) is a devastating condition, often resulting in long-term cognitive deficits. Not only does TBI present a major public health challenge, with approximately 50 million people worldwide effected each year (Maas et al., 2017), these injuries increase the risk of dementia and disability later in life (Mendez, 2017). Further, TBI has a significant economic burden, with a projected global cost of $400 billion US annually (Maas et al., 2017). As there are no effective treatments for TBI and 5.3 million people living with a TBI-related disability in the United States alone (Langlois and Sattin, 2005), there is a considerable unmet clinical need for novel therapeutics.

Neural stem cell (NSC) transplants have multiple potential mechanisms of action (MOA), including replacement of damaged cells (Haus et al., 2016), modulating the inflammatory microenvironment (Pluchino et al., 2003), and/or providing trophic support (Weston and Sun, 2018; Willis et al., 2020). With multifaceted potential, transplantation of NSCs may be a promising treatment strategy for TBI. Efficacy has been reported using human fetal NSCs (Wennersten et al., 2004; Gao et al., 2006), embryonic stem cell (ESC)-derived NSCs (Haus et al., 2016; Beretta et al., 2017) and induced-pluripotent stem cell (iPSC)-derived NSCs (Wei et al., 2016; Furmanski et al., 2019). As all these studies have applied cells from on-going culture, there is a significant gap in our understanding of the efficacy of freshly thawed cells from cryobanked stocks of hNSCs – a more clinically-relevant source for TBI.

The importance of assessing freshly thawed cells is highlighted by the mesenchymal stromal cell (MSC) clinical trials that failed translation from fresh, small-scale preparations in animal studies to frozen aliquots for humans (Galipeau and Sensébé, 2018). Further, continuous cell culture is an expensive, inefficient and time-consuming manufacturing strategy, presenting a major barrier to large-scale clinical deployment (Lipsitz et al., 2016; Badner et al., 2017). Additionally, as cell behavior (e.g., growth kinetics, secreted factors, expression of genes) is susceptible to physicochemical parameters (pH, dissolved oxygen, temperature, concentrations of exogenous factors), cell product purity and reproducibility remains problematic in continuous culture paradigms (Lipsitz et al., 2016). Production of cryopreserved cells provides a suitable alternative, as it allows for banking large quantities, storage until time of need, and shipment to diverse patient sites (Woods et al., 2016). Despite the potential, cryopreservation and freshly thawed cells present limitations. There have been reports of reduced immunomodulatory, blood regulatory properties and altered biodistribution directly after thawing compared to cells obtained freshly from culture (Chinnadurai et al., 2016; Moll et al., 2014; François et al., 2012). Nevertheless, to our knowledge, there have been no studies assessing the efficacy of freshly thawed hNSCs. Therefore, we aimed to test the therapeutic potential of our previously validated Shef-6 human embryonic stem cell (hESC)-derived NSC product (Haus et al., 2016; Beretta et al., 2017) following long-term cryostorage and thawing prior to transplant.

Additionally, long-term survival of hNSCs in xenotransplantation has been problematic, and frequently requires the use of immunodeficient rodents and dual immunosuppression (Anderson et al., 2011). We added anti-Asialo GM1 antibody injections to block natural killer cells (Kasai et al., 1981) in an effort to increase long-term hNSC survival.

Finally, this project investigates novel potential mechanisms of action of hNSCs for TBI. We provide evidence that freshly thawed Shef-6.0 hESC-derived NSCs (Shef 6.0 F/T) engraft within endogenous neurogenic niches, the meninges and lining of lateral ventricles, potentially through SDF-1alpha/CXCR4 ligand/chemokine receptor-mediated migration. We also show significant cognitive improvements in spatial learning and memory, anxiety, accompanied by ipsilateral host hippocampus sparing, evidence of rescued/improved neurogenesis, and a reduction in neuroinflammation. With anti-Asialo GM1 antibody, we report 53.4% cell survival, compared to 8.7% (Haus et al., 2016) and 3.9% (Beretta et al., 2017) found in our previous studies without supplemental immunosuppression Overall, these findings validate the potential of long-term bio-banking of hNSCs for TBI and demonstrate that cognitive improvement may stem from a trophic and/or anti-inflammatory response in endogenous neurogenic niches.

## Materials and Methods

### Experimental Design

All experiments, including animal housing conditions, surgical procedures, and postoperative care, were in accordance with the Institutional Animal Care and Use Committee guidelines at the University of California–Irvine (UCI). The use of human cells was approved by UCI’s Human Stem Cell Review Oversight Committee. In the first cohort, twenty-one male athymic nude (ATN) rats (NCI RNU/568, 50 days old, Charles River Laboratories, San Diego, CA) were randomly divided into three experimental groups. Due to surgical complications at time of transplant, two animals were excluded from the study and the final animal numbers were: sham (n=6), TBI+vehicle (n=7), and TBI+Shef6.0 F/T hNSCs (n=6). A second cohort of 18 animals, n=6 per group, was generated for cerebrospinal fluid (CSF) collection from the cisterna magna at two weeks post-cell transplant. Figure 1 provides an overview of experimental timeline and animal cohorts.

**Figure 1.**
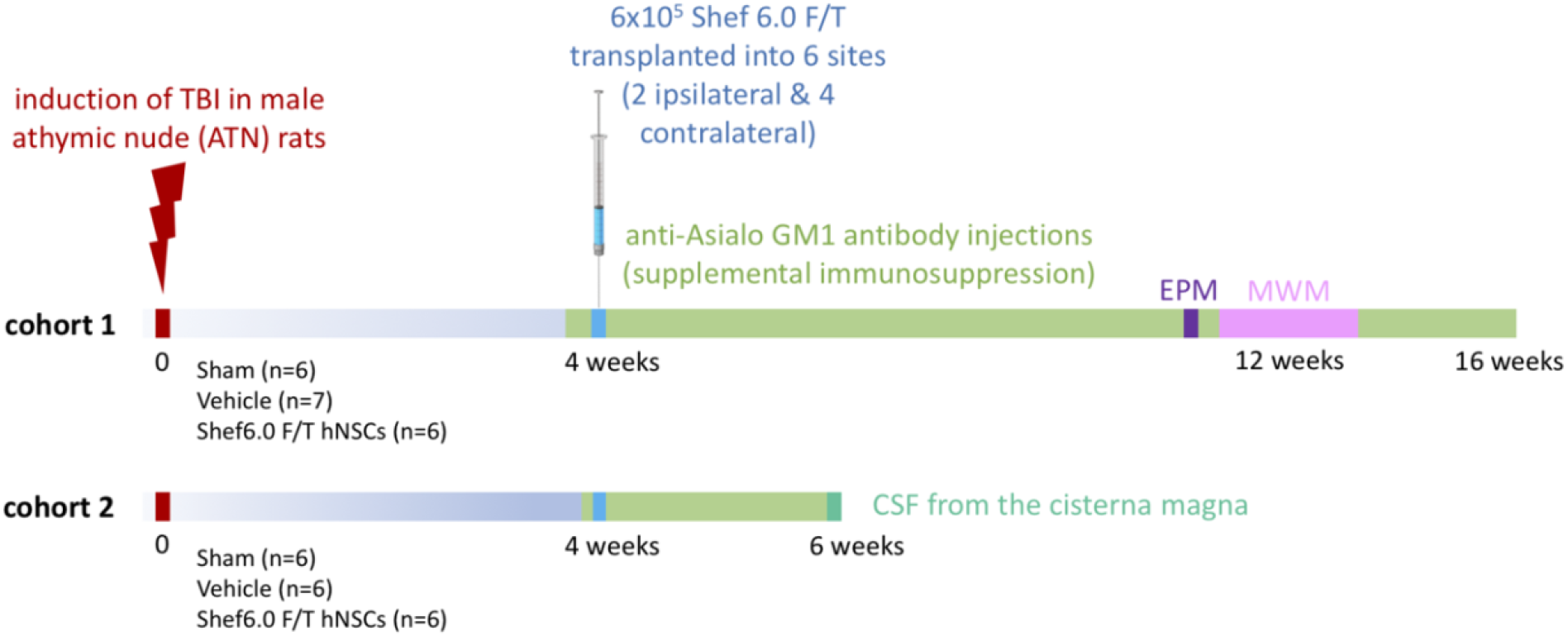
Experimental design and timeline. The study was comprised of two animal cohorts with male athymic nude (ATN) rats that underwent controlled cortical impact device-mediated unilateral traumatic brain injuries (TBI) and transplant of freshly thawed (F/T) human Shef 6.0 embryonic stem cell (ESC)-derived neural stem cells (NSCs) at 4 weeks post-injury. The animals were randomly divided into three experimental groups, where the first cohort had: sham (n=6), vehicle (n=7), and Shef6.0 F/T hNSCs (n=6). These animals were used for neurobehavioral testing, including elevated plus maze (EPM) and Morris water maze (MWM), as well as histological assessment at 16 weeks post-injury. The second cohort, n=6 per group, was generated only for cerebrospinal fluid (CSF) collection from the cisterna magna at two weeks post-cell transplant (6 weeks post-injury).

### Controlled Cortical Impact (CCI) Traumatic Brain Injury (TBI)

Animals were anesthetized with 3% isoflurane and placed in a stereotaxic frame (Leica Microsystems Inc). For the craniotomy (centered at A/P −4.5, M/L −3.6 relative to bregma), a small divot was made on the ipsilateral side (A/P −0.80, M/L −1.5) to serve as the reference point for the other two drill sites on transplant day. After making a small divot at the center of the craniotomy, a trephine was used to make a 6 mm burr hole, exposing the brain with an intact dura. Both divots were created using a Dremel^®^ rotary tool (Racine, WI; set at 5000 rpm). The angle of the stereotaxic was rotated to 15 degrees to make the craniotomy plane flat, before moving to a pneumatic controlled cortical impact device (TBI 0310; Precision Systems and Instrumentation, Fairfax Station, VA). The depth of the impact (5 mm flat metal impactor tip) was set at 2.5 mm with a velocity of 4.5 m/s and a dwell time of 500 ms. TBI coordinates were: A/P − 4.5 mm, M/L − 3.6 mm (left motor cortex, centered over the hippocampus). Immediately after procedure, a skull cap was glued to protect the brain lesion, the wound closed and animals received 10 ml of Lactated Ringer’s (50 mL/kg) subcutaneously (for a total of 3 days), 0.6 ml of buprenorphine (0.5 mg/kg), one dose subcutaneously immediately after surgery, then once in the AM and once in the PM the following day, and lastly, once more in the AM the second day after procedure, and 0.2 ml of enrofloxacin (2.5 mg/kg) subcutaneously immediately after surgery and thereafter daily for 5 days.

### Preparation and Transplantation of Shef6.0 F/T Human Neural Stem Cells (hNSCs) at 4 weeks Post-TBI

Shef6.0 hNSCs were derived from Shef6.0 hESCs (University of Sheffield, UK) in accordance with Human Stem Cell Research Oversight (hSCRO) and Institutional Biosafety Committee (IBC) protocols at UCI (Haus et al., 2014). The CD133^+^/CD34^−^ Magnetic-Activated Cell Sorted (MACS) Shef6.0 hNSCs were frozen in CryoStem ACF Freezing Medium (Biological Industries, CAT#05-710-1D) following monolayer culture at passage 27 at a density of 5.2 × 10^6^ cells. Following long-term (more than 6 months) storage in an ultralow temperature cryofreezer at −150°C (with gas phase liquid nitrogen backup), the cells were thawed the day of transplantation in a water bath at a steady 37°C, then transferred to a 15mL conical tube with 8mL pre-warmed Xeno-Free Neural Stem Cell Media (XF-NSM) consisting of X-Vivo Basal Medium (Lonza, 04-744Q), 1X N2 Supplement (Invitrogen, 17502-048), 10ng/mL LIF (Millipore, LIF1010), 100 ng/mL bFGF (Invitrogen, PHG0026), and 100 ng/mL EGF (Invitrogen, PHG0311). The cells were centrifuged at 0.2 × 1000 rcf for 5 minutes and all but 1mL of the supernatant was aspirated. A cell count via Hemocytometer was performed to confirm density. Following cell counts and determination of percent viability, the cells were then resuspended in XF-NSM to obtain a concentration of 100,000 cells/μL in a total volume of 25μL.

Each day, the cells were aliquoted to a specific random colored cap of a 1.5mL screw top gasketed microcentrifuge tube. The tube was screwed onto the cap which acted as the ‘lid’ and cells were delivered to the surgery suite for transplantation along with the vehicle samples (XF-NSM aliquoted at 25μL in different colored caps than the cells). The surgical team was blind to the sample color code, which varied by day. The percent viability of cells was calculated for the three different samples per day over two transplant days (at time of thaw and end of day). For the first animal cohort, cell viability was as follows: Day 1 (time of thaw): 93% (sample 1), 96.6% (sample 2) and 97.9% (sample 3). Day 1 (end of day, 8-10 hours post-thaw): 88% (valued for pooled samples). Day 2 (time of thaw): 73% (sample 1), 90.8% (sample 2) and 87.9% (sample 3). Day 2 (end of day, 8-10 hours post-thaw): 68.1% viability (valued for pooled samples).

For transplantation (at 4 weeks post-TBI), the animals were anesthetized with 3% isoflurane and placed in a stereotaxic frame. A Dremel^®^ rotary tool (Racine, WI), set at 5,000 rpm, was used to create 1 ipsilateral and 2 contralateral injection burr holes. A 10 μL Hamilton syringe (Cat#87930; Hamilton Company, Reno, NV) with a 1” 30g blunt needle was mounted into an UMP-3 (World Precision Instruments, Sarasota, FL) injector connected to a SYS-Micro4 controller (both from World Precision Instruments, Sarasota, FL). Subsequently, 6 injections of vehicle solution (XF-NSM) or 600,000 Shef6.0 F/T hNSCs in XF-NSM were made at the following coordinates relative to bregma (ipsilateral injection #1 A/P −0.8, M/L −1.5, D/V −2.8; ipsilateral injection #2: A/P −0.8, M/L −1.5, D/V −1.5; contralateral injection #3 A/P −2.5, M/L +2.5, D/V −4.9; contralateral injection #4 A/P −2.5, M/L +2.5, D/V −2.5; contralateral injection #5 A/P −5.3, M/L +4.4, D/V −4.7; contralateral injection #6 A/P −5.3, M/L +4.4, D/V −3.0). Prior to transplant, cells or vehicle were first triturated 3 times with a 10-μL pipette, then pulled up into syringe at a rate of 417 nl over 1 minute. For each injection site, the needle was initially lowered an additional 0.10 mm to create a pocket in the brain. For each of the six 1μl injections, a dose of 100,000 cells/μL over 2 minutes was administered with a dwell time of 2 min after the second injection of each burr hole. The total dose was 600,000 hNSCs. Bone wax was applied to seal each of the burr holes. The incision was closed with wound clips and post-operative care was administered as for CCI surgeries.

### Anti-Asialo GM1 Antibody Injections (Supplemental Immunosuppression)

Rabbit anti-asialo GM1 antibody (Wako Chemicals, 986-10001) was reconstituted (1mg) in milliQ water (1mL) and serile saline (9mL). Subsequently, a 0.5mL (0.5ug dose) was delivered to each experimental animal via intraperitoneal injection at −1day, +14day, +35day, +56day and +77day, relative to the transplantation day.

### Collection of Cerebrospinal Fluid (CSF) from the Cisterna Magna

At two weeks following cell transplant surgeries in the second animal cohort, each rat (n=6 per group) was anesthetized in a 3% isoflurane chamber, shaved and cleaned with Betadine solution. The animals were subsequently secured with ear bars in a stereotaxic frame while being maintained on a heating pad and an anesthesia mask delivering 3% isoflurane. The head was angled downward at approximately 45° to expose the dorsal neck region. A medial incision of the skin was made along the occipital ridge directly above the atlanto-occipital (a-o) membrane, as previously described (Pegg et al., 2010). The three neck and shoulder muscle layers were separated to expose the a-o membrane, which was thoroughly cleaned using a cotton swab to allow clear access to the cisterna-magna; this reduces blood contamination during CSF collection. A 27g butterfly needle fixed to a polyethylene line and syringe was carefully inserted through the membrane into the cisterna-magna up to the end of the needle bevel (no more than 1-2 mm). While the needle was held in place, a second surgeon slowly drew ~100-200 μL of CSF into the line or until the appearance of blood was observed, at which point the tubing was clamped and cut. The sample without blood contamination was then stored at −80°C for later cytokine analysis. The muscle layers were sutured, the incision was closed with wound clips, and post-operative care was administered as for CCI surgeries.

### Cytokine Array for CSF Proteomic Profiling

The R&D Systems rat cytokine XL ELISA Proteome Profiler array (ARY30, R&D Systems Inc., Minneapolis, MN, https://www.rndsystems.com) was used as per manufacturer’s instructions with minor modifications. CSF from each condition (n=6 per group) was randomized, run on a single proteome array membrane, and analyzed by an individual blinded to the group identity. Briefly, Alexa Fluor™ Streptavidin-conjugated 488 (Catalog number S32354, ThermoFisher Scientific) was used at a 1:500 dilution (30 minutes, room temperature) as replacement for the kit’s Streptavidin‐horseradish peroxidase to increase detection sensitivity. Proteome array membranes were scanned on an Azure Biosystems C600 gel/blot imager. The scans were quantified using Western Vision’s HLImage++ Analysis software. The vehicle control and Shef6.0 F/T-treated sample values were normalized to sham CSF and expressed as fold change.

### Shef6.0 F/T hNSCs Transwell Migration Assay

The migration assay was completed as previously described (Hooshmand et al., 2017). Briefly, 150 μL of differentiation media with varying concentrations of recombinant human SDF1alpha (Peprotech, Catalog 300-28A) was added to the wells of the feeder tray (Millipore), and 100 μl dissociated Shef6.0 hNSC (300,000 cells/ml) was added to the migration chambers, followed by incubation for 3h at 37°C. Migration chambers were removed, placed on new 96-well trays containing 150 μl prewarmed cell detachment buffer in the wells, and incubated for 30 min at 37°C. At the end of the incubation, 50 μl 1:75 dilution of CyQuant GR Dye:Lysis buffer was added to the cell detachment buffer and incubated for 15 min at room temperature. Finally, 150 μl CyQuant GR Dye:Lysis/detachment solution was transferred to a new 96-well plate, and migration was quantified using a 480/520-nm filter set on a fluorescent plate reader. The experiment was conducted in biological triplicate.

### Cognitive Behavioral Testing

EthoVision XT version 13 software (Noldus, Leesburg, VA) was used to video record and track animals during cognitive behavioral testing. Behavioral assessment began 8 weeks post-transplantation (12 weeks post-injury). Experimenters were blinded to the experimental assignment of the animals. Animals were handled daily by all experimenters for at least one week prior to the start of behavioral assessment.

### Elevated Plus Maze (EPM) for Risk-Taking Behavior

The EPM consisted of two open arms and two closed arms (width of 10cm and length of 112cm across two arms), made of black plastic (Med-Associates, Fairfax, VT). The test was performed in a dark room and a white noise machine (57dBa ± 1 dBa) was used to mask extraneous sounds outside of the testing room. Infrared beam detection and an infrared video camera suspended above the maze facilitated video recording and animal tracking during the test. The walls and floors of the maze were cleaned with 70% ethanol at the start of each testing day and between each animal. On test day, animals are placed at the center point of the EPM and allowed to roam freely in the maze for 5 minutes. Cumulative time spent in the open arms was used as the primary measure of risk-taking. Distance travelled and velocity were also assessed for each animal, to validate the presence/absence of confounding differences in motor function.

### Morris Water Maze (MWM) for Spatial Learning and Memory

The Morris water maze (MWM) used a circular fiberglass pool (185cm diameter) filled with water maintained at 25.0 ± 0.5°C and non-toxic white paint was added to make the water opaque. An acrylic platform (10cm × 10cm) was placed 1-2cm below the surface of the water. Distinct spatial cues were attached to the walls of the testing room. A video camera was centered above the pool and video output was processed via a computer in an adjoining testing room (Windows 10 Pro, 1024 × 768, 25 fps). MWM testing was conducted with the room lights on and a white noise machine (57dBa ± 1 dBa) was used to mask extraneous sounds outside of the testing room. The MWM test consisted of five phases: cued pre-training (day 0), acquisition (days 1-5), probe test (day 6), reversal (days 7-11) and reversal probe test (day 12). During cued pre-training (day 0), a flag was placed on the hidden platform to mark its location in the pool, and the spatial cues on the walls were removed. Animals had four 60-second trials to explore the pool and swim to the cued platform. If they did not find and sit on the platform by the end of the trial, they were directed to the platform by the experimenter. The platform was moved to one of the four quadrants of the pool for each of the four trials (NE, SE, SW, NW).

During the remaining MWM phases (days 1-12), the flag marking the platform location was removed and the spatial cues were returned to the walls of the testing room. The acquisition phase (days 1-5) was used to assess the animals’ spatial learning and consisted of four 60-second trials during each of the five days. During this phase, the platform was positioned in the SW quadrant of the pool. The animals were placed into the water for each trial, facing the pool wall, at one of four different starting positions (NW, N, E, SE) chosen in semi-random order. They were allowed to swim freely within the pool until 1) they landed on the platform and remained for three seconds or 2) the 60-second trial ended. If the animal did not land on the platform during the trial, they were directed to the platform by the experimenter. The latency to land on the platform was recorded for each trial. The probe test (day 7) was used to assess their spatial memory of the platform location and consisted of just one 60-second trial. During this test, the platform was removed from the pool and the animals were placed into the water at a novel starting location (NE). Their swim path and time spent in platform quadrant (SW) were recorded.

The reversal phase (days 8-12) measured the animals’ ability to learn a new location of the platform, assessing their learning flexibility. The protocol for this phase was the same as the protocol for acquisition, with the exception of the platform location (NE) and the starting positions for each trial (SE, S, W, NW). The latency to land on the platform was recorded for each trial. Finally, the reversal probe test (day 13) assessed the animals’ spatial memory of this new platform location. The protocol for this test was the same as the previous probe test, except that animals were placed into the water at a different starting location (SW). The animals’ swim paths and times spent in the platform quadrant (NE) were recorded.

### Automated Water Maze Analysis (Rtrack)

Analysis of spatial exploration data from the MWM task was completed using *Rtrack*, an open-source software package (https://cran.r-project.org/web/packages/Rtrack/index.html).

### Tissue Processing

At 12 weeks post-transplantation (16 weeks post-injury), animals were sacrificed for histological assessment as previously described (Haus et al., 2016) with minor modifications. Briefly, animals were anesthetized with a lethal dose of Fatal-Plus pentobarbital sodium (100 mg/kg, i.p.) and transcardially perfused with 100 mL of phosphate-buffered saline (PBS), followed by 300-400 mL of 4% paraformaldehyde (PFA). Brains were dissected and post-fixed in 4% PFA and 20% sucrose at 4°C for 48 hours. Tissue was then flash frozen at −65°C in isopentane (2-methyl butane) and stored at −80°C. For cryosectioning, sets of four frozen brains containing all treatment groups were embedded in NEG-50 (6502, Thermo Fisher Scientific, Waltham, MA). Brains were sectioned coronally in 30μm thick sections using a cryostat and mounted using a CryoJane tape transfer system (Leica Biosystems, Inc., Buffalo Grove, IL).

### Histological Analysis

#### Nissl Stain

Sections were stained with Cresyl violet, as previously described (Gold et al., 2018), for quantification of corpus callosum, lateral ventricle, lesion and total brain volumes. In short, blinded experimenters drew contours around the structures in question on every 12^th^ section of tissue, using StereoInvestigator software v11.08.01 (MicroBrightField, Inc., Williston, VT). A 300 μm grid was overlaid to assess corpus callosum, lateral ventricle, lesion and total brain volume with a counting frame of 30 μm; these parameters yielded a coefficient of error (CE) for each animal that was less than 0.10.

### 3,3’-Diaminobenzidine (DAB) Immunohistochemistry for STEM121 Cell Survival Analysis

All immunohistochemistry was conducted at room temperature following antigen retrieval with Buffer A (Electron Microscopy Sciences, 62706-10). For 3,3’-diaminobenzidine (DAB) immunohistochemistry, sections were washed in 0.1M Tris, incubated in 3% hydrogen peroxide/10% methanol for 30 minutes, and washed again in 0.1M Tris. Sections were then permeabilized in a 0.1% Triton X-100 wash for 15 minutes and then blocked for one hour in bovine serum albumin (BSA) and normal serum from the species in which the secondaries were raised. Sections were then incubated in primary antibody, STEM121 (human cytoplasm, 1:2000; Y40410, Takara, Mountain View, CA), overnight at room temperature to detect engrafted cells. The next day, sections were incubated with a biotin-conjugated, purified IgG *anti-*mouse secondary antibody (1:500; 715-066-151, Jackson ImmunoResearch, West Grove, PA) pre-adsorbed against the species in which the primary was raised in, followed by the avidin-biotinylated peroxidase complex (ABC) using the Vectastain Elite ABC kit (Vector Laboratories, USA) and prepared according to the manufacturer’s recommendations. After several washes, the signal was visualized with diaminobenzidine (DAB) (Vector Laboratories, USA). Sections were counterstained with hematoxylin, dehydrated, and coverslipped using DEPEX mounting media (Electron Microscopy Sciences, Hatfield, PA).

### Stereology for STEM121 Cell Survival Analysis

Histological sections, at 30 μm, comprising the entire rostral-caudal region of the cortex (A/P + 5.16 to A/P − 9.36) were used to estimate total human cell survival. Unbiased stereological analysis was performed using the Microbrightfield Stereoinvestigator (MBF Bioscience) Optical Fractionator probe by an observer blind to treatment group. A section interval of 1/12 was utilized, placing adjacent sections analyzed at 360 μm apart. Contours were drawn at the 4× objective, outlining the full brain, and human cell identification as well as counting was performed at 63× magnification. The counting frame utilized was 150 × 150 μm with a grid size of 500 × 500 μm. The dissector height was set at 12 μm with an upper and lower guard zone of 1 μm.

### Fluorescent Immunohistochemistry

Sections were washed in PBS following antigen retrieval and then permeabilized in a 0.1% Triton X-100 wash for 15 minutes. Sections were then blocked for one hour in PBS with 0.1% Triton X-100 and normal serum from the species in which the secondaries were raised, and then incubated in appropriate primary antibodies overnight at room temperature. Primary antibodies used were: polyclonal rabbit *anti-*Olig2 (1:500; AB9610, Millipore, Burlington, MA), polyclonal goat *anti*-CXCR4 (1:100; AB1670, Abcam, Cambridge, MA), polyclonal rabbit *anti*-SDF1alpha (1:100, AB9797, Abcam, Cambridge, MA), polyclonal rabbit *anti-*Iba1 (1:1000; 019-19741, Wako, Osaka, Japan), monoclonal mouse *anti-*STEM121 (1:500; Y40410, Takara, Mountain View, CA), polyclonal rabbit *anti-*GFAP (1:500; Z0334, Dako, Agilent, Santa Clara, CA), polyclonal goat *anti*-DCX (1:250; sc-8066, Santa Cruz Biotechnology, Dallas, TX), polyclonal rabbit *anti*-NeuN (1:500; ABN78, Millipore, Burlington, MA), polyclonal rabbit *anti-*AQP1 (1:1000; AB2219, Millipore, Burlington, MA). The next day, sections were washed three times (5 minutes each), blocked in normal serum from the species the secondary antibodies were raised in for one hour, and then incubated in secondary antibodies for two hours, followed by Hoechst 33342 counterstain (1:500; H1399, Life Technologies, Thermo Fisher Scientific, Waltham, MA). The secondaries used were: AlexaFluor 488-conjugated F(ab’)2 fragment donkey *anti*-goat (1:500; A11055, Invitrogen, Thermo Fisher Scientific, Waltham, MA), AlexaFluor 555-conjugated F(ab’)2 fragment donkey *anti*-rabbit (1:500; A31572, Invitrogen, Thermo Fisher Scientific, Waltham, MA), AlexaFluor 647-conjugated F(ab’)2 fragment donkey *anti*-mouse (1:500; A31571, Life Technologies, Thermo Fisher Scientific, Waltham, MA), and AlexaFluor 555-conjugated F(ab’)2 fragment goat *anti*-rabbit (1:500; A21428, Invitrogen, Thermo Fisher Scientific, Waltham, MA). After five 5-minute washes with PBS, sections were coverslipped with Fluoromount-G^®^ (SouthernBiotech, Birmingham, AL) mounting media, sealed with nail polish, and allowed to dry overnight. Imaging was performed on an Olympus FV3000 Confocal Laser Scanning Microscope (Olympus, Tokyo, Japan). Laser percentage, voltage, gain, and offset were kept constant during imaging for each quantification parameter using a secondary only control.

### Imaris Quantification (Iba1+ and DCX+ Cells)

Iba1+ cells were counted by blinded personnel using the spot detection and batch features of IMARIS-based 3D analysis software (Bitplane, Zurich, Switzerland). DCX morphology was assessed using the filament branching feature of IMARIS to quantify the number of cell projections within a set region of interest, specifically the dentate gyrus. IMARIS was also used to quantify the number of DCX+ nuclei within the subgranular zone of the dentate gyrus.

### Statistical Analysis

Prior to statistical analysis, and blind to treatment group, a Grubbs’ test was performed to identify potential outliers from each group (α = .05). For some single-time point assessments (eg. chemotaxis assay), unpaired t-tests or one-way analyses of variance (ANOVA) were performed with Tukey or Dunnett’s post hoc test, as specified. Other behavioral analyses employed, when specified, (non)linear mixed-effects models in R and MATLAB to capture changes over time (i.e., trial, day). Post hoc tests in R were performed using its *glht* function and post hoc tests in MATLAB were performed using its *coefTest* function. Statistical significance was set at *p* < .05. Error bars in figures are SEM. Unless otherwise stated, statistics were computed using GraphPad Prism (GraphPad Software Inc., La Jolla, CA).

## Resultss

### Long-Term Engraftment of Transplanted Shef 6.0 F/T hNSCs in the Meninges and the Lining of Lateral Ventricular Wall – Most Remain as Neural Progenitors

In accordance with the experimental overview (Figure 1), Shef 6.0 F/T hNSCs survival and engraftment was assessed at 12 weeks post-transplantation (16 weeks post-injury) using an Optical Fractionator probe. Although many cells were located within the brain parenchyma, the majority were observed in the meninges (Figure 2A) and lining the lateral ventricular wall (near the subventricular zone, SVZ; Figure 2A). Unbiased stereological assessment found that of the 600,000 cells transplanted an average of 53.4% survived at three months post-transplantation. In contrast, our previous studies using Shef 6-derived hNSCs without the use of anti-asialo GM1 antibody, had cell survival at 8.7% and 3.8% (Haus et al., 2016; Beretta et al., 2017, respectively). However, given the different quantification parameters used between these studies, the increase in cell survival presented in here may not solely result from the application of anti-asialo GM1 antibody. Importantly, of the engrafted Shef 6.0 F/T cells (Figure 2B-C), 19.5 ± 4.7% expressed NeuN, 19.7 ± 3.2% expressed markers of astrocytes or glial progenitors (GFAP), and 14.2 ± 2.2% expressed markers of early oligodendrocytes (Olig2). Despite some expression of more mature neuronal lineage markers, many of the transplanted cells (78.1 ± 2.3%) also expressed doublecortin (DCX), a marker of immature neurons or neural progenitors. Differentiation profiles are shown for human cells within the meninges and those lining the lateral ventricular wall (Figure 2D).

**Figure 2.**
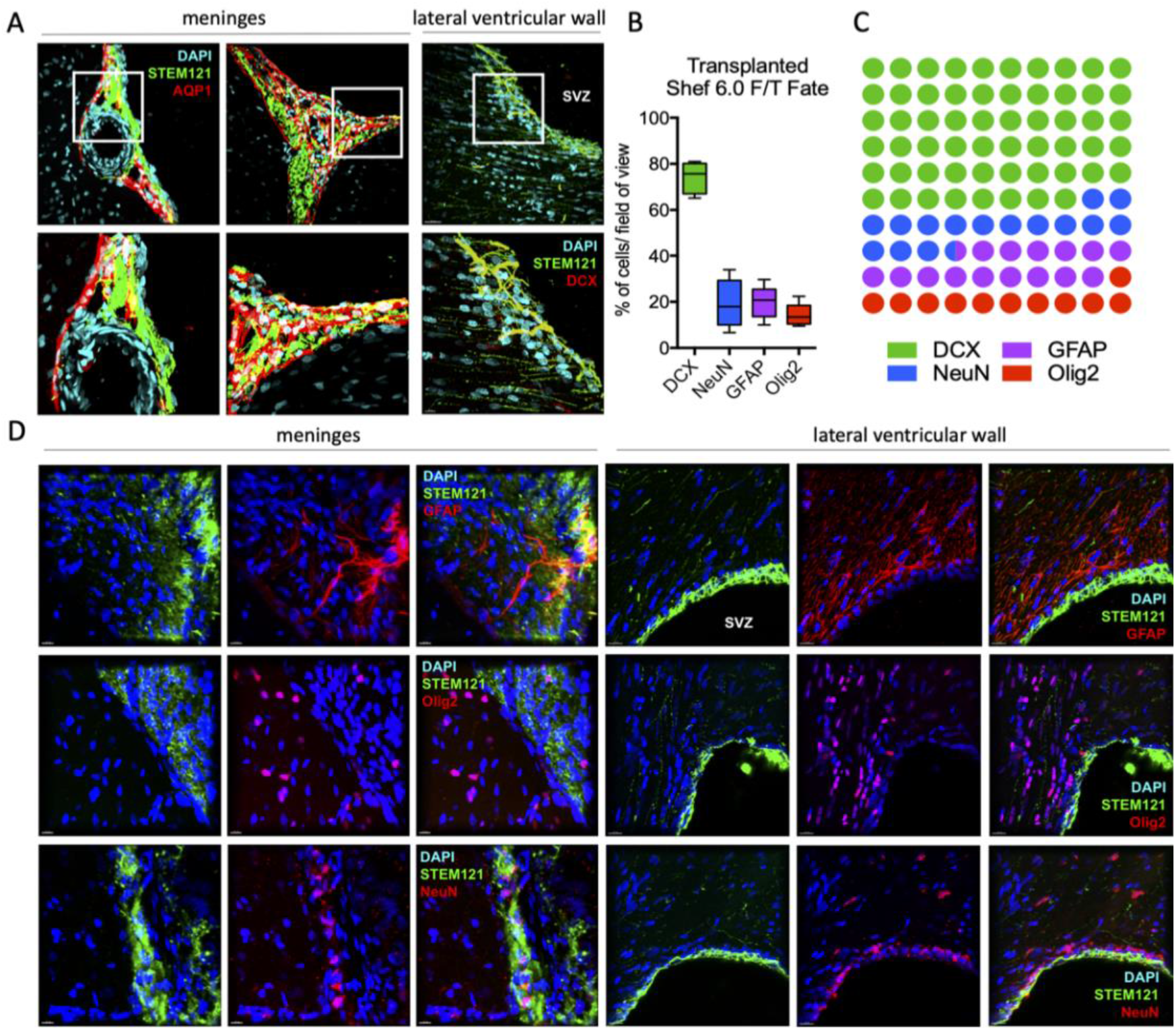
Freshly thawed ESC-derived Shef 6.0 hNSCs display long-term engraftment in the meninges and the lining of the lateral ventricular wall with tri-lineage potential. **A,** A majority of transplanted Shef 6.0 hNSCs (human cytoplasm, STEM121) were found in the meninges and the lining of the lateral ventricular wall (near the subventricular zone, SVZ) at three months post-transplantation. **B,** Quantification (*n*=5) of engrafted Shef 6.0 hNSC fate proportions indicate that the majority of cells remain as doublecortin positive neural progenitors. Data are expressed as median ± SEM. **C,** While transplanted cells differentiated into neurons (NeuN), astrocytes (GFAP) and oligodendrocytes (Olig2), many remained doublecortin-expressing progenitors (DCX). **D,** Representative images of differentiated Shef 6.0 hNSCs in the meninges and lining of the lateral ventricular wall. All images were taken at 40X on the Olympus FV3000 laser-scanning confocal spectral inverted microscope. Scale bar=20μm.

### CXCR4 Expression and SDF1alpha-driven Shef 6.0 F/T hNSCs Migration Could Account for Meningeal as well as Lateral Ventricular Wall Engraftment

In *vitro*, Shef 6.0 F/T hNSCs express chemokine receptor CXCR4 (Figure 3A) and display a significant, *F*(3, 20) = 9.909, *p* = .0003 (One-way ANOVA, Dunnett’s multiple comparison post hoc), chemotactic response to SDF1alpha at 100-200ng (Figure 3B). Further, at three months post-transplantation, Shef 6.0 F/T hNSCs maintain expression of CXCR4 and appear to engraft in SDF1alpha-rich regions of the meninges (Figure 3C) and the lining of the lateral ventricular wall (near the subventricular zone, SVZ; Figure 3D).

**Figure 3.**
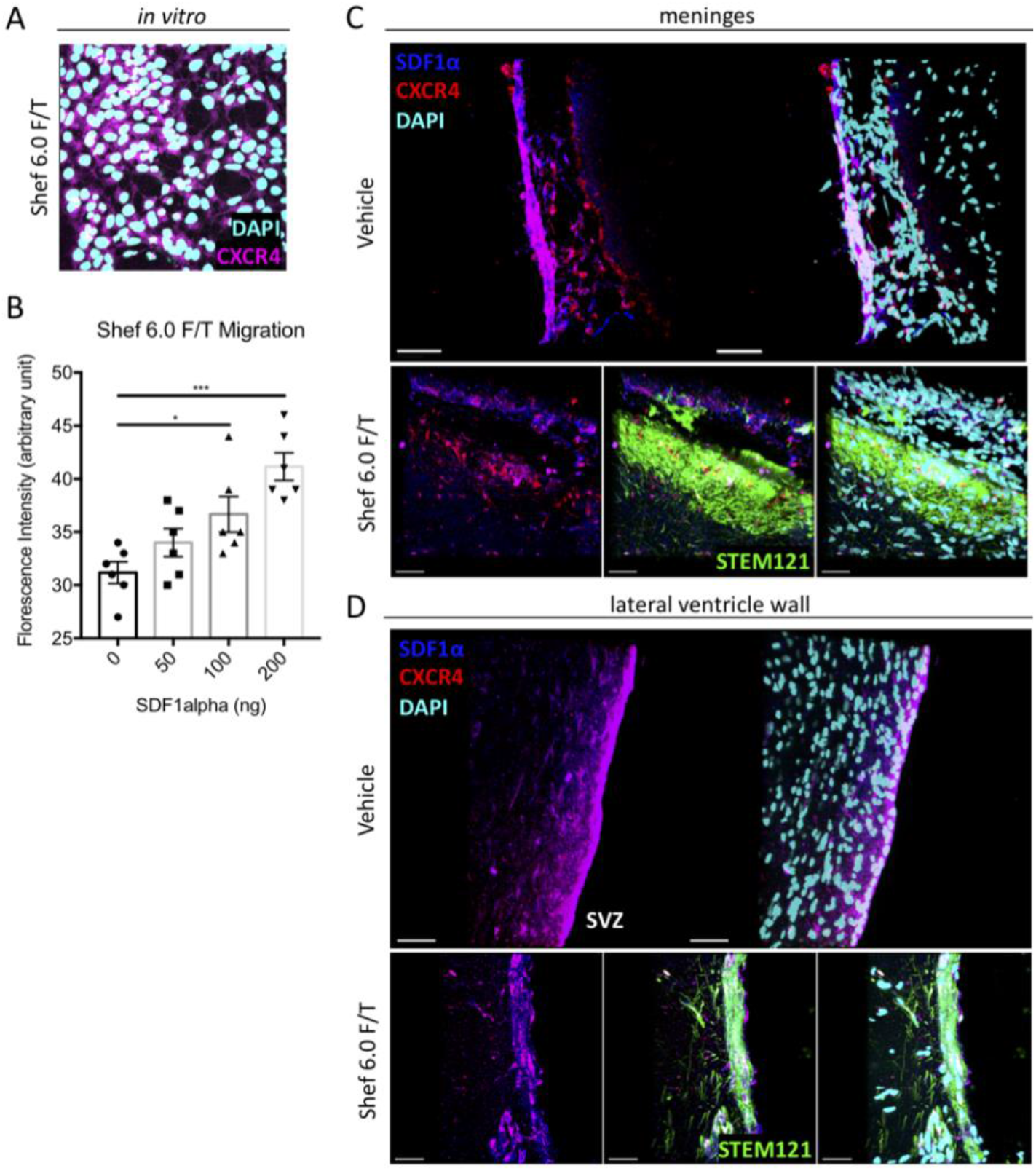
Freshly thawed ESC-derived Shef 6.0 hNSCs express CXCR4 and SDF1 alpha-driven cell migration. **A,** Representative image of CXCR4 expression *in vitro* in Shef 6.0 hNSCs. **B,** A three-hour migration assay shows Shef 6.0 hNSCs are responsive to SDF1 alpha-induced chemotaxis. Data are expressed as mean ± SEM. One-Way ANOVA, with Tukey multiple comparisons post hoc; *, p ≤ .05; s***, p≤ .001. **C-D**,Representative images of the meninges (C) and the lateral ventricular wall (D) three months post-transplant further highlight that Shef 6.0F/T hNSCs (green) maintain CXCR4 expression *in vivo* (red) and engraft in regions rich in SDF1 alpha (dark blue). Arrows indicate multiple STEM121 immunopositive hNSCs colocalized with CXCR4 and SDF1alpha. Images were taken at 40X (Vehicle) and 60X (Shef6.0F/T) oil objectives on an Olympus FV3000 laser-scanning confocal spectral inverted microscope. Scale bar=20μm.

### Chronic Transplantation of Shef 6.0 F/T hNSCs Did Not Resolve Tissue Loss nor Ventricular Swelling Following TBI

Gross tissue anatomical changes (Figure S1) were analyzed following injury and Shef 6.0 F/T transplantation via stereology using a Cavalieri estimator. Specifically, total brain volume (Figure S1A) remained unchanged, *F*(2, 15) = 1.046, *p* = .3755 (One-way ANOVA, Tukey’s multiple comparison post hoc) following injury in vehicle (3.40×10^11^ ± 2.3 ×10^10^ μm^3^) or Shef 6.0F/T transplanted (3.80×10^11^ ± 2.1 ×10^10^ μm^3^) animals in comparison to shams (3.44×10^11^ ± 3.6 ×10^10^ μm^3^). As expected, the ipsilateral lateral ventricle was significantly enlarged, *F*(2, 15) = 3.753, *p*=.04 (One-way ANOVA, Tukey’s multiple comparison post hoc) and corpus callosum volume was reduced, *F*(2, 15) = 5.695, *p*=.02 (One-way ANOVA, Tukey’s multiple comparison post hoc; Figure S1B-C) in injured animals in comparison to shams. No differences were found in Shef 6.0 F/T-treated rat ventricle as well as corpus callosum volume (2.2×10^10^ ± 4.3×10^9^ μm^3^ for ipsilateral lateral ventricle, 2.0×10^10^ ± 1.6×10^9^ μm^3^ for corpus callosum) compared to that of vehicle-treated rats (2.6×10^10^ ± 4.8×10^9^ μm^3^ for ipsilateral lateral ventricle, 1.9×10^10^ ± 1.4×10^9^ μm^3^ for corpus callosum). Similarly, Shef 6.0 F/T transplantation (1.5×10^10^ ± 3.2×10^9^ μm^3^) did not alter lesion volume, *F*(6, 5) = 2.827, *p* = .2740 (unpaired t-test) following injury when compared to vehicle (2.1×10^10^ ± 4.9×10^9^ μm^3^) treated injured controls (Figure S1D). Three-dimensional representative images illustrate that the volumetric outcomes were not grossly different between the vehicle and cell-treated animals (Figure S1E). While TBI caused lateral ventricle enlargement, corpus callosum reduction, and a persistent lesion cavity following injury, chronic transplantation of Shef 6.0 F/T did not resolve any gross tissue loss, ventricular swelling, corpus callosum volume, nor lesion volume.

### Chronic Transplantation of Shef 6.0 F/T hNSCs Improved Spatial Learning and Memory on the MWM and reduced risk-taking behavior on the EPM

Spatial learning and memory were assessed using Morris water maze (MWM) at 8 weeks post-transplantation (12 weeks post-injury) following transplantation of Shef 6.0 F/T or vehicle controls (Figure 4). Latency to platform, analyzed separately for acquisition and reversal, was assessed via nonlinear mixed-effects models using the *nlme* function in R (Figure 4B). The fitted nonlinear function was an exponential decay function [A*((1-k)^(Trial-1))], in which *A* was the intercept of the function and *k* was the decay rate. Here, *k* is equivalent to learning rate (i.e., higher learning rates reflect faster times to find the platform). The nonlinear mixed-effects model included both by-group fixed effects and by-rat random effects on *A* and *k*. For acquisition, analysis revealed main effects of group on learning rate, *F*(2, 356) = 6.47, *p* = .002, but not intercept, *F*(2, 356) = 1.28, *p* = .280. Post hoc tests revealed that the Sham rats exhibited faster learning rates than both the Vehicle and Shef 6.0 rats, *p*s ≤ .003, while the Vehicle and Shef 6.0 rats were not significantly different, *p* = .594. Similarly, during reversal, there was no main effect of group on intercept, *F*(2, 356) = 0.07, *p* = .934, but there was a main effect of group on reversal learning rate, *F*(2, 356) = 13.42, *p* < .001. Here, while the Sham rats showed much faster reversal learning than Vehicle rats and Shef 6.0 rats, *p*s < .001, Shef 6.0 rats showed significantly faster reversal learning than Vehicle rats, *p* = .018.

**Figure 4.**
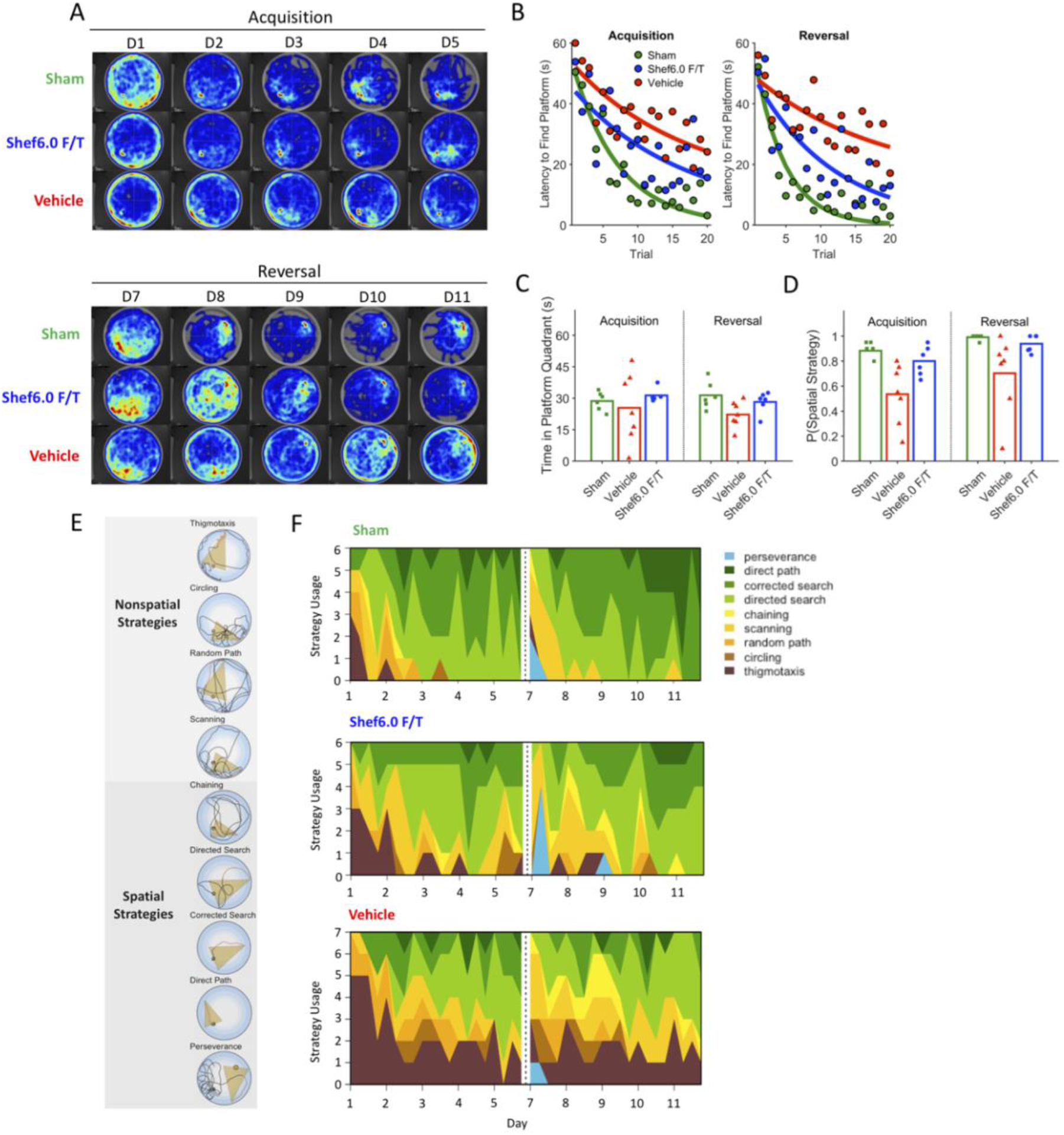
Chronic transplantation of freshly thawed ESC-derived Shef 6.0 hNSCs improved spatial memory and learning following traumatic brain injury. **A**, Heatmap images showing the cumulative time spent within the maze during the acquisition (week 8 post-transplantation) and reversal phases (week 9 post-transplantation). **B,** Morris water maze (MWM) latency to platform curves and, **C**, time spent in the platform quadrant on probe test days, in which the platform was removed from the pool. With respect to the former, data points reflect group means in each trial, and the curvilinear functions are exponential decay functions, in which steeper functions reflect faster learning of platform location (i.e., faster times to find the platform). On reversal learning, Sham rats exhibited faster reversal learning than Vehicle rats and Shef 6.0 hNSC rats, but Shef 6.0 hNSC rats showed significantly faster learning than Vehicle rats on a nonlinear mixed-effects model (*nlme* in R). For the probe-test figures, bars reflect group means and the data points are individual rats. Further analysis was conducted on the likelihood of using a spatially dependent strategy to locate the hidden platform during training trials. **D**, Shams were significantly more likely than the vehicle group to spend time in the platform quadrant during the probe test, whereas the Shef6.0 hNSC group was not significantly different from Shams; *, p ≤ .05; **, p ≤ .01. **E,** Examples of spatial and non-spatial MWM search strategies recognized by the classification algorithm used (*Rtrack*). **F,** The number of rats that uses these respective strategies within each group on the MWM task is shown, with each strategy color-coded.

For the probe test days, in which the hidden platform was removed from the pool, we analyzed the amount of time (out of the 60-s probe trial) spent in the quadrant of the pool that had contained the platform during training (Figure 4C). Linear models with a between-subjects factor of group were separately conducted on acquisition and reversal data. Analysis was performed in MATLAB. There were no significant differences between groups in acquisition, *F*(2, 16) = 0.51, *p* = .609, but there was a main effect of group in reversal, *F*(2, 16) = 4.06, *p* = .037. With respect to reversal, the sham group was significantly more likely than the vehicle group to spend time in the platform quadrant during the probe test, *p* = .013. In contrast, the Shef6.0 F/T group was not significantly different from the sham, *p* = .358, or vehicle group, *p* = .090.

Further assessment of the platform search strategies classified search strategies (Garthe et al., 2009) as either being spatially or non-spatially dependent (Figure 4E), highlighting differences between groups during the acquisition and reversal phases (Figure 4F). Generalized linear mixed-effects models (binomial distribution, logit link; Figure 4D) were used to compare the use of spatial search strategies between Shef 6.0 F/T-treated, vehicle controls, and sham injury animals. The criterion was whether animals used a spatial strategy (i.e., scanning, chaining, directed search, corrected search, direct path; coded as 1) or not (i.e., thigmotaxis, circling, random path; coded as 0). Perseverance-type trials were not included in analysis. Each phase (acquisition, reversal) was analyzed separately. Analysis was performed in MATLAB. The models included a fixed effect of group and by-rat random intercepts. Intuitively, this analysis assessed the probability of each group using spatial versus nonspatial strategies to find the hidden platform during acquisition and reversal. In acquisition, there was a main effect of group, *F*(2, 377) = 9.17, *p* < .001, in which the sham and Shef6.0 F/T groups were significantly more likely to use spatial search strategies than vehicle rats, *p*s ≤ .004. In contrast, there was not a statistically significant difference between sham and Shef6.0 F/T groups with respect to the likelihood of spatially dependent search-strategy usage, *p* = .214. Similarly, there was a main effect of group in reversal, *F*(2, 367) = 5.13, *p* = .006. As in acquisition, while there was no significant difference between sham and Shef6.0 F/T groups, *p* = .181, these two groups were significantly more likely to use spatially dependent strategies during the reversal phase than the vehicle group, *ps* ≤ .028.

The Elevated Plus Maze (EPM) was used to assess risk-taking behavior (Figure 5). There was a significant reduction (One-way ANOVA, Tukey’s multiple comparisons post hoc) in the time spent in open arms following transplantation of Shef 6.0 F/T compared to vehicle treated TBI controls (*F*(2, 16) = 10.49, *p* = .0012; Figure S2D), demonstrating reduced risk-taking among the Shef 6.0 F/T group. These results were not confounded by any differences in distance travelled (Figure 5B) or velocity (Figure 5C).

**Figure 5.**
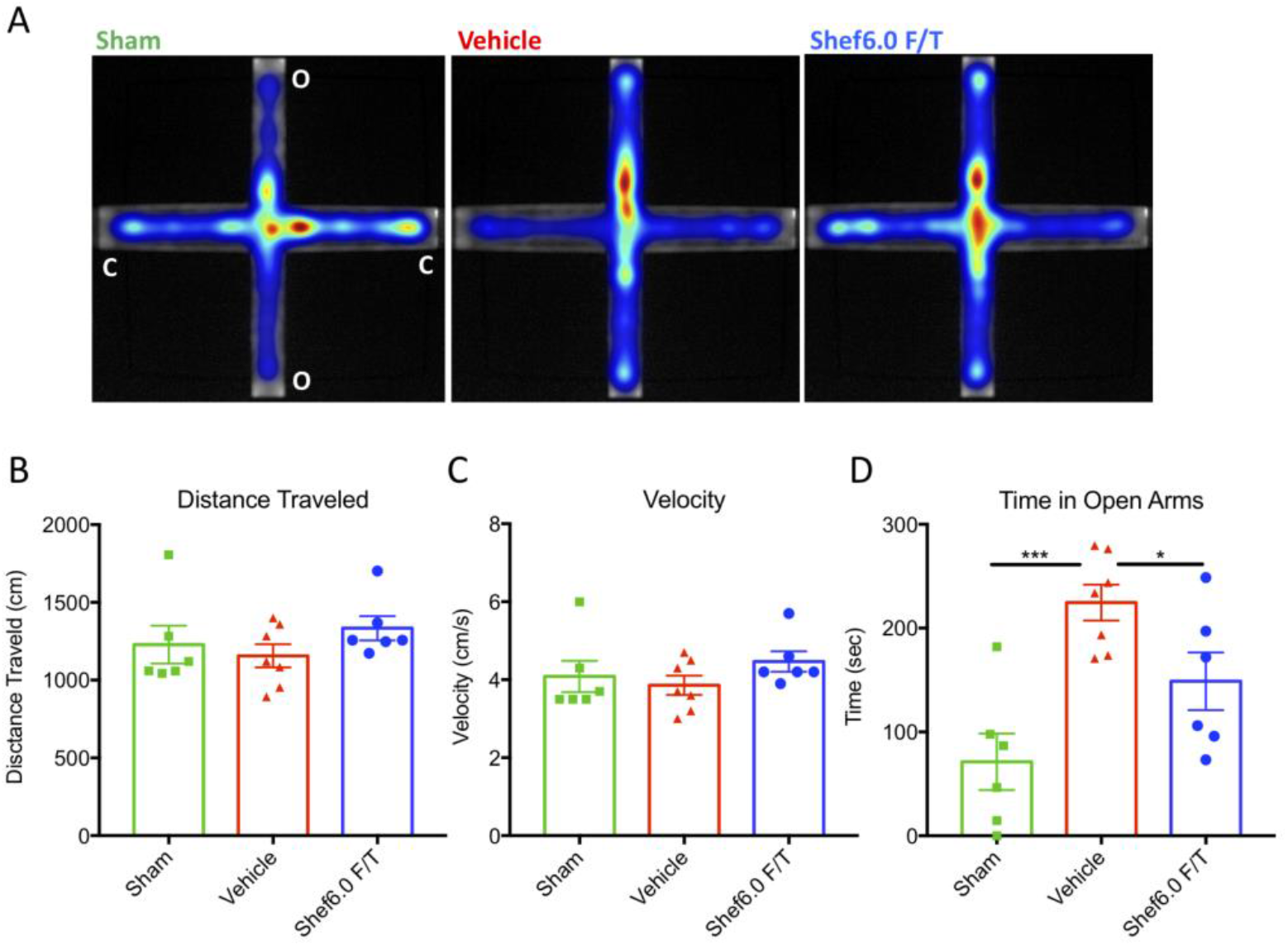
Chronic transplantation of freshly thawed ESC-derived Shef 6.0 NSCs reduced risk-taking behavior post-TBI. **A,** Heat maps showing the cumulative time spent in the open (o) and closed (c) arms of the elevated plus maze complete at week 8 post-transplantation. Heat maps are generated to show the cumulative behavior of all experimental animals per group. The total (*n* = 6-7 per group), **B**, distance travelled, **C**, velocity and, **D**, time in open arms are shown. Data are expressed as mean ± SEM. One-Way ANOVA, with Tukey multiple comparisons post hoc; *, p ≤ .05; ***, p < 0.001.

### Shef 6.0 F/T Transplantation Improves Host Hippocampal Neuron Survival and Progenitor Morphology

Considering the aforementioned improvements in learning and memory and reduction in risk-taking following Shef 6.0 F/T transplantation, hippocampal neuron survival was assessed ipsilateral and contralateral to the injury. Specifically, an Optical Fractionator probe was used to quantify neuronal changes in the cornu ammonis (CA) and dentate gyrus (DG). The number of pyramidal CA neurons was significantly reduced after injury compared to sham on the ipsilateral side (Figure 6A; sham: 5.9×10^5^ ± 9.7×10^4^; vehicle: 1×10^5^ ± 9.7×10^4^; *p* = .003; Shef 6.0 F/T: 3×10^5^ ± 3.5×10^4^; *p* = .008, One-way ANOVA, Tukey’s multiple comparison post hoc) as well as in the CA3 on the contralateral side (Figure 6A; sham: 5×10^5^ ± 1.4×10^4^; vehicle: 3×10^5^ ± 4.8×10^4^; p = .01, One-way ANOVA, Tukey’s multiple comparison post hoc). There were also significantly more ipsilateral CA neurons following Shef 6.0 F/T transplantation compared to the vehicle control (p = 0.04, One-way ANOVA, Tukey’s multiple comparison post hoc), without an effect on the contralateral CA3 (Figure 6A). Similarly, the number of DG neurons was also significantly reduced after injury on the ipsilateral side (Figure 6B; sham: 1.1×10^6^ ± 1.2×10^5^; vehicle: 4×10^5^ ± 5.4×10^4^; *p* = .0007; Shef 6.0 F/T: 5×10^5^ ± 8.7×10^4^; *p* = .004, One-way ANOVA, Tukey’s multiple comparison post hoc), but without differences on the contralateral side. Although there were no differences in ipsilateral or contralateral DG neurons between Shef 6.0 F/T-treated rats and vehicle controls (Figure 6B), contralateral DG volume was found to be trending towards greater (*p* = .06 One-way ANOVA, Tukey’s multiple comparison post hoc) following Shef 6.0 F/T transplantation (Figure 6C). Further analysis of contralateral DG changes revealed a trending increase (p = .08, Figure 6E, One-way ANOVA, Tukey’s multiple comparison post hoc) in the number of DCX+ cell projections through the DG relative to the number of DCX+ cell bodies in the subgranular zone (SGZ). Representative images are shown in Figure 6D.

**Figure 6.**
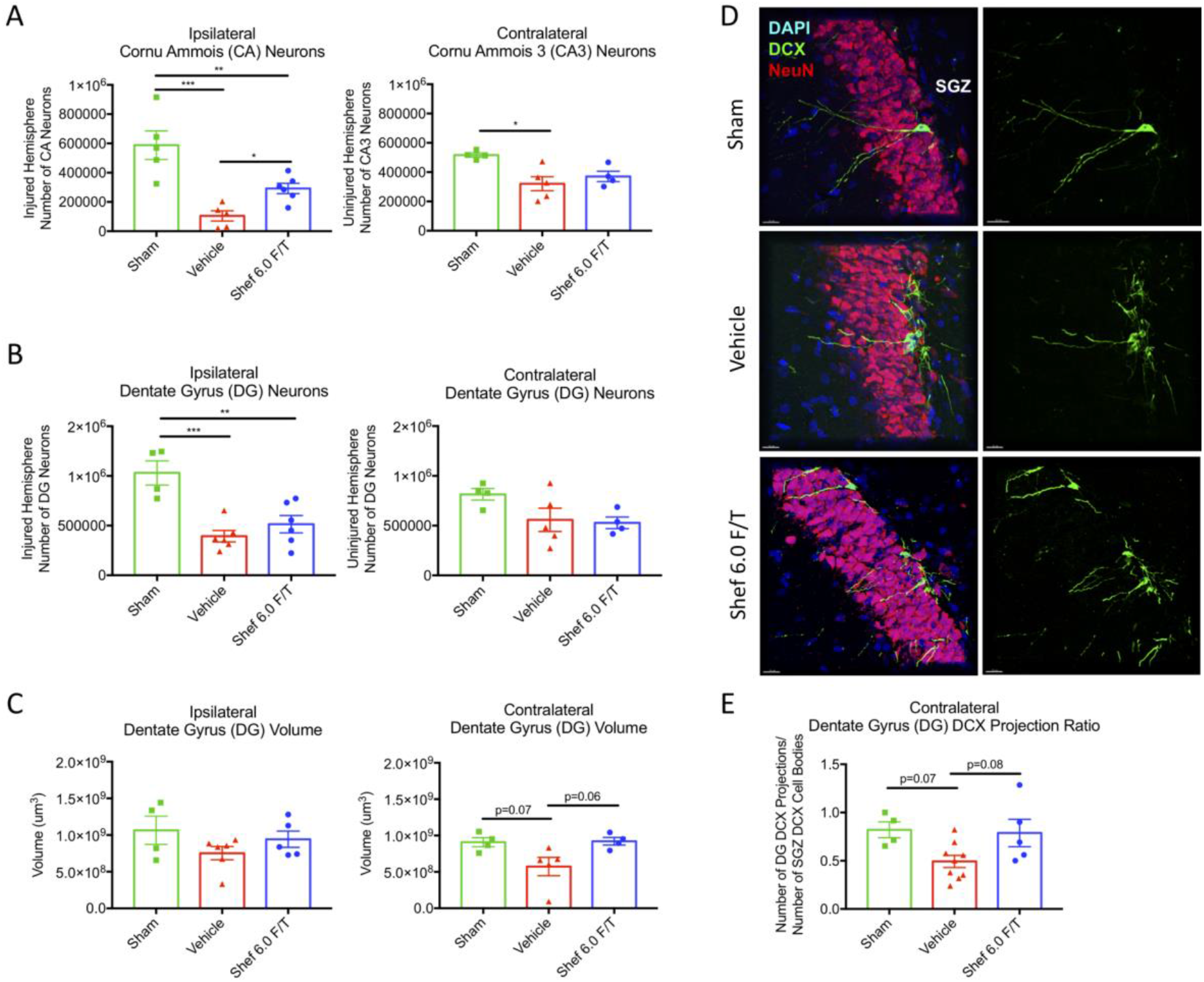
Chronic transplantation of freshly thawed ESC-derived Shef 6.0 hNSCs preserved contralateral cornu ammois hippocampal integrity and improved ipsilateral hippocampal progenitor morphology post-TBI. **A,** Stereological assessment of host cell numbers revealed that Shef 6.0 F/T hNSC transplantation increased preservation of ipsilateral cornu ammois (CA) neurons without any significant changes to contralateral CA3 neurons. **B,** Ipsilateral dentate gyrus (DG) neuron numbers were reduced following injury without any rescue of host cells following Shef 6.0 F/T hNSC transplantation. **C,** A Cavalieri estimator of volume suggested that Shef 6.0 F/T hNSC transplantation led to a trend in increased contralateral DG volume via with no difference on the ipsilateral hemisphere. **D,** Representative images of DCX+ neural progenitors and their projections in the subgranular zone (SGZ) of the contralateral DG are shown for each condition. Images were taken at 63X on the Leica TCS SP8 confocal microscope. The ratio of projections into the dentate per cell body is indicative of successful integration into the hippocampal circuit. **E,** There was a trending increase in doublecortin (DCX) projection to cell body ratio in the Shef 6.0 hNSC group contrasted to the vehicle group. Data are expressed as mean ± SEM. One-Way ANOVA, with Tukey multiple comparisons post hoc; *, p ≤ .05; **, p ≤ .01; ***, p ≤ .001.

### Chronic Shef 6.0 F/T hNSC Transplantation Reduced Neuroinflammation Following TBI

Cerebral spinal fluid was collected from the cisterna magna at two weeks following Shef 6.0 F/T hNSC transplantation (Figure 7A) and profiled using the R&D Systems rat XL cytokine array kit (ARY030). A representative heat map of the proteome profile was generated using the BROAD Institute’s R implementation of Morpheus with Euclidean distance hierarchical clustering (Figure 7B). Sham CSF was initially compared with samples from vehicle controls, where discovery was determined using the two-stage linear step-up procedure of Benjamini, Krieger and Yekutieli, with Q = 1%. Of the 80 cytokines/chemokines screened, 43 discoveries were identified (CCL17/TARC, Adiponectin/Acrp30, Hepassocin, MMP-2, RAGE, Prolactin, IL-2, IL-6, Cyr61/CCN1, WISP-1/CCN4, CCL2/JE/MCP-1, MAG/Siglec-4a, SCF, CXCL7/Thymus Chemokine-1, IL-17A, IGF-I, IL-3, MMP-3, CX3CL1/Fractalkine, Neprilysin/CD10, Fibulin 3, IL-13, Resistin, NT-3, CCL3/CCL4/MIP-1a/ß, HGF, FGF-21, MMP-9, FGF acidic, IFN, IL-1ra/IL-1F3, GM-CSF, IL-4, ICAM-1/CD54, CXCL2/GROß/MIP-2/CINC-3, FGF-7/KGF, CCL11/Eotaxin, Serpin E1/PAI-1, Galectin-1, NT-4, Flt-3 Ligand, IL-22, CCL20/MIP-3a). Of these, t-tests with Holm-Sidak correction for multiple comparisons identified 6 cytokines (RAGE, MMP-2, IL-2, CCL11/Eotaxin, Adiponectin/Acrp30 and Hepassocin) to be significantly (*ps* < .01) different between the vehicle and Shef6.0F/T-treated animals.

**Figure 7.**
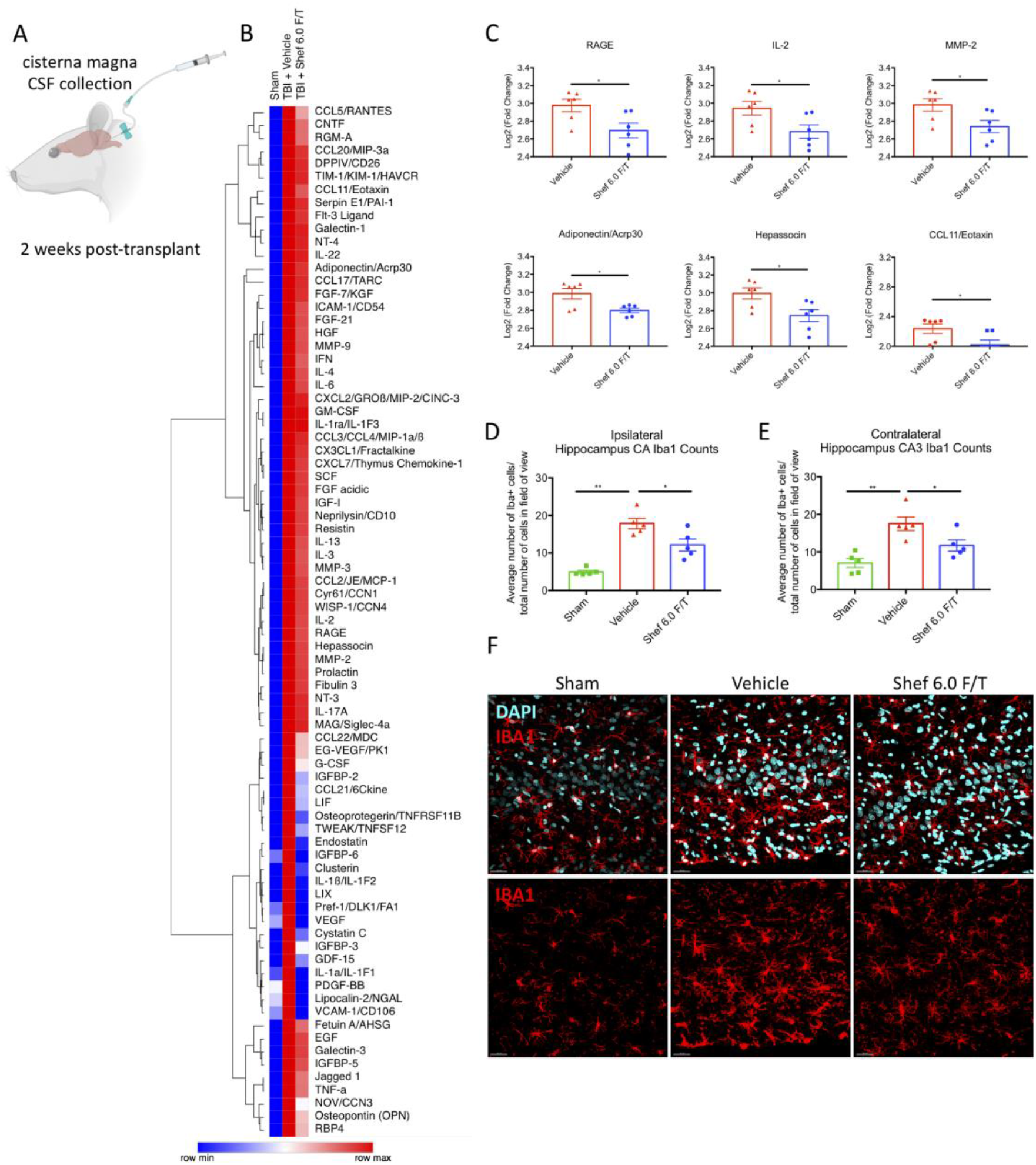
Chronic transplantation of freshly thawed ESC-derived Shef 6.0 hNSCs reduced inflammatory cytokines in cerebrospinal fluid and the number of hippocampal microglia post-TBI. **A,** Cerebrospinal fluid (CSF) was collected from the cisterna magna (image generated via BioRender) at two weeks following Shef 6.0F/T hNSC transplantation and profiled using the R&D Systems rat XL cytokine array kit (ARY030). **B**, A representative heat map was generated using the BROAD Institute’s R implementation of Morpheus with Euclidean distance hierarchical clustering. **C,** Data are expressed as mean Log2 (fold change) relative to sham CSF (*n* = 6 per group) and statistically significant differences in cytokine expression (multiple *t* tests with Holm-Sidak correction for multiple comparisons) are shown for RAGE, IL-2, MMP-2, Acrp30, Hepassocin, and CCL11. **D,** To measure potential effects on the microglial response post-injury, we counted the number of Iba1 immunopositive cells within the hippocampus. The number of ipsilateral hippocampal cornu ammois (CA) microglia was increased following traumatic brain injury and significantly reduced following Shef 6.0 F/T hNSC transplantation. **E,** The contralateral CA3 also had reduced microglia (Iba1) following Shef 6.0 F/T transplantation. Data in D and E are expressed as mean ± SEM. One-Way ANOVA, with Tukey multiple comparisons post hoc; *, p ≤ .05; **, p≤ .01. **F,** Representative CA3 images from the contralateral hippocampus are shown for each condition.

Hippocampal neuroinflammation was assessed via microglia/macrophage (Iba1) counts ipsilateral (Figure 7D) and contralateral (Figure 7E) to the injury. As expected, there was a significant rise in ipsilateral, *F*(2, 12) = 26.66, *p* < 0.0001 (One-way ANOVA, Tukey’s multiple comparison post hoc) and contralateral, *F*(2, 12) = 11.91, *p* = .001, (One-way ANOVA, Tukey’s multiple comparison post hoc) cornu ammonis (CA) microglia/macrophage numbers following TBI. Further, Shef 6.0 F/T transplantation reduced the number of Iba1 positive cells both ipsilateral (*p* = .018) and contralateral (*p* = .047) to the injured hemisphere. To contextualize, Shef 6.0 F/T transplantation resulted in a 32.1% decrease in ipsilateral and 33.1% decrease in contralateral Iba1 positive microglia/macrophages.

## Discussion

We report the first successful application of freshly thawed hNSCs from cyrobanked stocks for chronic TBI. In addition to cognitive improvements in the MWM and EPM following hNSC transplantation post-TBI, which validate previous findings (Haus et al., 2016; Beretta et al., 2017), we present novel evidence of long-term hNSC engraftment within endogenous neurogenic niches, changes in developing immature host neurons in the dentate gyrus (DG), and reductions in neuroinflammation. We also demonstrate that administration of anti-asialo-GM1 increased transplanted human cell survival to 53.4%, compared to 8.7% (Haus et al., 2016) and 3.9% (Beretta et al., 2017) found in previous studies without supplemental immunosuppression. These results highlight the clinical feasibility of freshly thawed Shef-6.0 human ESC-derived NSCs for chronic TBI and provide insight into multiple putative mechanisms of action (MOA).

Expanding on studies involving sub-acute (nine days post-injury) transplantation of hNSCs post-TBI in ATN rats (Haus et al., 2016; Beretta et al., 2017), this work extends the potential therapeutic window by transplanting freshly thawed hNSCs (Shef-6.0 F/T) at 30 days post-injury, at a more clinically relevant chronic phase of TBI (Leo and McCrea, 2016). However, the spatial distribution and differentiation of engrafted NSCs in the injured brain proved surprising after chronic transplantation. Many of the cells were engrafted in the lining of the lateral ventricular wall, near the subventricular zone (SVZ), and within the meninges (Figure 2A), at sites known to maintain populations of endogenous NSCs (Lois and Alvarez-Buylla, 1993; Decimo et al., 2011; Bifari et al., 2017; Nakagomi and Matsuyama, 2017). Re-examination of tissue from Haus 2016 and Beretta 2017 also revealed the presence of transplanted hNSCs within these niches. Although meningeal and ventricular engraftment was unexpected, NSCs have been previously reported to home to the neurovascular niche in a stromal-derived factor 1alpha (SDF1alpha) and CXC chemokine receptor 4 (CXCR4) dependent manner (Kokovay et al., 2010). Consistently with this pathway, Shef-6.0 hNSCs were found to express CXCR4 (Figure 3A) and exhibited a significant chemotactic response to SDF1alpha at 100-200ng *in vitro* (Figure 3B). A knock-out of CXCR4 in hNSCs followed by loss of migration or *in vivo* transplantation would further validate this interaction.

Even at three months post-transplant, 78% of transplanted cells expressed doublecortin (DCX), a marker of neuronal precursor cells and immature neurons. While some cells also differentiated into NeuN+ cells, GFAP+ cells, and Olig2+ cells, the lasting expression of DCX may stem from the unique engraftment profile within known neurogenic niches of the adult brain. Previous work where SVZ-derived rat NSCs were transplant into the hippocampus reported that many cells retained “stemness” (DCX expression) even at three months after transplant (Shetty and Hattiangady, 2016); we would expect human cells to take longer to mature than rodent cells.

The modest proportion of differentiation among engrafted cells suggests that integration within host tissue is not the only, or even primary, mechanism underlying cognitive improvements. Consistent with earlier studies, Shef-6.0 hNSCs were able to significantly improve MWM reversal learning (Figure 4B), a well-established hippocampal-dependent learning task. Although time spent in the probe quadrant during acquisition as well as reversal was not significant, the Shef-6.0 hNSC times during the reversal probe test were sandwiched in between sham and vehicle values, indicative of partial recovery (Figure 4C). Classifying search behaviors and associated swim patterns using a parameter-based algorithm (Overall et al., 2020; Garthe et al., 2009; Figure 4E), sham and Shef6.0 hNSC groups were significantly more likely to use spatial strategies than vehicle controls during both acquisition and reversal (Figure 4D). A breakdown of the unique navigation strategies highlights these group differences, which were most apparent during reversal (Figure 4F). The shift towards more effective and directed strategies has been uniquely attributed to adult-generated granule cells within the DG that become selectively recruited into existing hippocampal networks (Kee et al., 2007; Garthe et al., 2009).

As the hippocampus is primarily involved in spatial learning and memory (Lassalle et al., 2000; Florian and Roullet, 2004), the significant effects on MWM navigation and EPM performance were associated with greater hippocampal integrity. We focused on the hippocampal dentate gyrus to cornu ammonis 3 (CA3) pathway known to be specifically involved in the acquisition of spatial memory (Lassalle et al., 2000; Florian and Roullet, 2004). Shef-6.0 F/T hNSCs significantly reduced the loss of host ipsilateral hippocampus CA neurons, largely damaged by the cortical contusion, without an effect on neuron numbers in the ipsilateral dentate gyrus (DG), where the damage was more extensive. Contralateral to the injury, while there was no significant difference between the vehicle or Shef-6.0 F/T treated CA or DG neurons, we did observe a trend towards greater DG volume (Figure 6C) with Shef 6.0 F/T-treated animals. The greater contralateral DG volume was also associated with changes in immature neuron (DCX+) morphology, where Shef-6.0 F/T treatment yielded a greater ratio of projections through the granular layer of the DG over number of subgranular zone cell bodies (Figure 6D-E). Injury-induced changes in the dendritic morphology of newborn hippocampal neurons has been previously reported in TBI (Villasana et al., 2015) and stroke (Niv et al., 2012; Woitke et al., 2017), and may indicative aberrant neurogenesis or maladaptive plasticity. Therefore, rather than, or in addition to the direct integration of transplanted cells, another MOA of hNSCs may be via the modulation of healthy host hippocampal neurogenesis or restoration of anatomically appropriate dendritic arborization. Validation of this possible MOA should be studied further, quantifying neurogenesis through bromodeoxyuridine labelling as well as dendritic reconstruction and quantification of synaptic redistribution post-TBI.

Aligned with the MWM results, hNSC treated animals displayed significantly reduced risk-taking behavior on the EPM (Figure 5D). While there may be a role of hippocampal glutamate in EPM (Xiang et al., 2011), which should be the subject of future study, there were no other gross neuroanatomical changes found with cell transplantation. Specifically, there were no histological differences in long-term brain, lateral ventricle, corpus callosum and lesion volumes between injury groups (Figure S1). The injury, and thereby lesion, has largely stabilized by 30 days post-injury, and was unchanged by cell transplantation. Together, this implicates the importance of hippocampal mediated MOAs via hNSCs.

We also characterized the effects of hNSCs on neuroinflammation, which persists for years following TBI (Plesnila, 2016). Specifically, microglia/macrophages can remain activated even several years after injury, as demonstrated in post-mortem human brain tissue (Johnson et al., 2013). Further, with a role in cell debris phagocytosis, antigen presentation (Hickey and Kimura, 1988), synapse pruning (Paolicelli et al., 2011) and dialogue with endogenous neural progenitors (Biber et al., 2007), lasting changes in microglia/macrophage numbers and activation state are indicative of tissue health (Donat et al., 2017). We observed a significant increase in hippocampal Iba1+ cells (microglia/macrophages) post-TBI (Figure 7D-F). Moreover, consistent with the preservation of hippocampal neurons and improvements in dentate neuron morphology, Shef-6.0 hNSCs significantly decreased the number of Iba1+ cells compared to controls (Figure 7D-F).

In tandem, we found a significant decrease in inflammatory cytokines in CSF collected two weeks following Shef-6.0 F/T transplantation in comparison to the vehicle group. When sham CSF was initially compared with samples from injured vehicle controls, 43 of 80 cytokines/chemokines screened via protein array were identified as significantly different, highlighting a persistent and lasting inflammatory response following injury. Importantly, of the 43 TBI-induced cytokines/chemokines, six were identified as significantly different between the vehicle and Shef-6.0 F/T treated animals (Figure 7C). Specifically, the levels of RAGE, MMP-2, IL-2, CCL11/Eotaxin, Adiponectin/Acrp30 and Hepassocin were reduced following Shef-6.0 F/T infusion. While the exact roles remain unknown, reductions in RAGE, MMP-2 and CCL11 have been reported to ameliorate neuroinflammation in a variety of models, including multiple sclerosis and Parkinson disease (Hannocks et al., 2019; Huang et al., 2020; Wang et al., 2020).

Importantly, as we report on a single timepoint in what is likely a dynamic and evolving process, the relationship between specific cytokines, the reduction in microglia, and changes in immature DG neuronal morphology, is just a snapshot. Nevertheless, together, these results suggest that Shef-6.0 F/T treatment may modulate neuroinflammation post-TBI via multiple MOAs. Future work should assess temporal cytokine changes as well as Iba1+ cell morphology/phenotype, which would provide greater insight to their polarization state and thereby their inflammatory role.

In summary, we demonstrate the feasibility of using freshly thawed hNSCs for the treatment of chronic TBI. Through a CXCR4-mediated chemotactic response to SDF1alpha, Shef-6.0 F/T were found to display long-term engraftment, primarily as DCX+ progenitors, in the meninges and in the lateral ventricular wall. Transplanted cells were associated with a reduction in inflammatory cytokines, a reduction in parenchymal inflammation, evidenced by fewer Iba1+ cells, and improved morphology and host integration of immature DG neurons. Consistent with hippocampal mediated effects, there was a significant improvement in spatial memory and learning, and a reduction in risk-taking behavior post-TBI. Using an automated parameter-based algorithm classification approach, the improvements in MWM navigation were further confirmed by a Shef-6.0 F/T specific increase the use of spatial search strategies. Overall, this new understanding of Shef6-derived hNSCs lays a foundation for future work and improvements in patient care.

## Acknowledgements

We want to thank technical staff at the Sue & Bill Gross Stem Cell Center, especially Rebecca Nishi, Christopher Nelson, Joshua David, and Krystal Carta for their help with animal surgeries, functional assessments and cell culture technical assistance. We are also particularly grateful for the generous support of Rupert W. Overall in sharing the Rtrack software package and assistance in the reproducible automation of water maze analysis. For video editing and quantitative analysis, we also thank students Ashlyn Wang, Mark Christian E. Marquez, Karan Bhatt and Deva Watson. We also thank Pranav Sankar for technical assistance with immunohistochemistry. This study was supported by CIRM DISC2-10195. Javi Lepe was a CIRM intern funded through the CIRM Grant EDUC2-08383.

---

**Figure S1.**
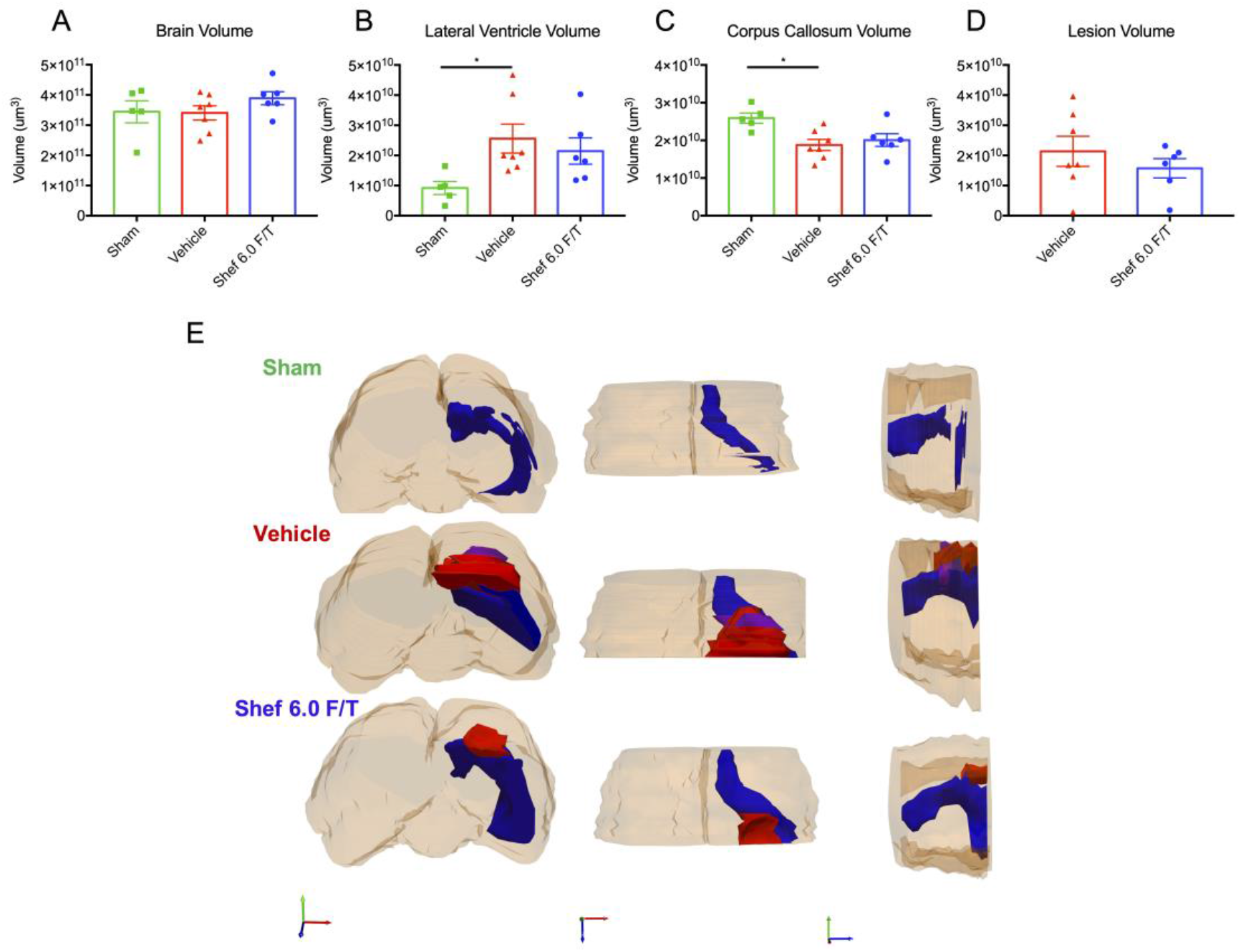
Stereological quantification of volume using the Cavalieri estimators for multiple regions. There were no changes to long-term, **A**, brain, **B**, lateral ventricle, **C**, corpus callosum and, **D**, lesion volumes following chronic transplantation of freshly thawed ESC-derived hNSCs (*n* = 6-7 per group). However, as expected, there was a significant increase in lateral ventricle volume and significant decrease in corpus callosum volume in the vehicle group in comparison to the shams. It is also important to note that neither ventricle volume nor corpus callosum volume were significantly different between the sham and Shef 6.0F/T hNSC treated animals. **E,** Three-dimensional representative renditions are shown for each condition, where the lesion is shown in red, lateral ventricle in blue and perspective overlap in purple. Data are expressed as mean ± SEM. One-Way ANOVA, with Tukey multiple comparisons post hoc; *, p ≤ .05.

